# Isolation and characterization of SARS-CoV-2 in Kenya

**DOI:** 10.1101/2022.08.22.504904

**Authors:** Albina Makio, Robinson Mugasiali Irekwa, Matthew Mutinda Munyao, Caroline Wangui Njoroge, Peter Kipkemboi Rotich, Tonny Teya Nyandwaro, Joanne Jepkemei Yego, Anne Wanjiru Mwangi, James Hungo Kimotho, Ronald Tanui, Vincent Rutto, Samson Muuo Nzou

## Abstract

The emergence of Severe Acute Respiratory Syndrome-Coronavirus-2 (SARS-CoV-2) from Wuhan, China, in December 2019 raised a global health concern that eventually became a pandemic affecting almost all countries worldwide. The respiratory disease has infected over 530 million people worldwide, with over 950,000 deaths recorded. This has led scientists to focus their efforts on understanding the virus to develop effective means to diagnose, treat, prevent, and control this pandemic. One of the areas of focus is isolation of this virus, which plays a crucial role in understanding the viral dynamics in the laboratory. In this study, we report the isolation and detection of locally circulating SARS-CoV-2 in Kenya. The isolates were cultured on Vero Cercopithecus cell line (CCL-81) cells, RNA extraction conducted from the supernatants, and reverse transcriptase-polymerase chain reaction (RT-PCR). Genome sequencing was done to profile the strains phylogenetically and identify novel and previously reported mutations. Vero CCL-81 cells were able to support the growth of SARS-CoV-2 in vitro, and mutations were detected from the two isolates sequenced (001 and 002). These virus isolates will be expanded and made available to the Kenya Ministry of Health and other research institutions to advance SARS-CoV-2 research in Kenya and the region.

**Author Summary:** The Coronavirus disease 2019 (COVID-19) pandemic is caused by a type of coronavirus that emerged in Wuhan, China in December 2019 and later spread to almost all countries. Many countries are still finding ways to contain it. The virus has been studied in many ways to investigate its origin, infectivity, and evolution. Different variants of the virus have emerged and spread, causing a lot of concern as to whether the pandemic will end soon. Significant studies have proven the ability of the virus to grow in the laboratory using cell lines that offer the necessary conditions. Therefore, this study sought to find out the growth of the virus in specific monkey cell line and the variant circulating within the Kenyan population. We found that the selected cell lines supported viral growth outside a human host system. In addition, the circulating virus was found to have evolved to enhance its survival mechanism. This is the first study in Kenya to report this virus’s isolation, culture, and identification in monkey kidney cells. These cells supported the growth of the virus in the laboratory and analysing the genome of the growth products showed the virus was related to previously reported strains with multiple changes in its whole DNA sequence.

## Introduction

The coronaviruses (*coronaviridae)* were first recognized as a new family of viruses in 1968. They are enveloped, positive-sense-stranded ribonucleic acid (RNA) viruses with a genome of 26-30kb, with the largest genome among all the known RNA viruses (1). The name of these viruses is derived from their morphology with spike proteins on their surface that appear like a crown shape (2). Phylogenetically, coronaviruses are classified into four genus: Alphacoronaviruses, Betacoronaviruses, Gammacoronaviruses, and Deltacoronaviruses. There are four human coronaviruses, namely, 229E, Netherland 63 (NL63), Organ culture 43 (OC43), and Hong Kong University 1(HKU1), which infect the upper respiratory tract and cause mild symptoms. There are three other coronaviruses of zoonotic origin that infect the lower respiratory tract of humans and can cause severe respiratory illness and lead to fatalities. These are severe acute respiratory syndrome coronavirus (SARS-CoV), Middle East respiratory syndrome coronavirus (MERS-CoV) (3) (4), and SARS-CoV-2.

However, recently, a new coronavirus strain emerged in Wuhan, China, that has caused a worldwide pandemic that was later named SARS-CoV-2 by the International Committee on Taxonomy of Viruses (5). Sequence and phylogenetic analysis of various published strains show close similarity (99% homology) to the wild type SARS-CoV strain (6). Research is still underway to understand the aetiology and pathophysiology of the virus.

As of 15^th^ July 2022, COVID-19 had infected over 560 million individuals with over 6 million deaths globally, of which 9.1 million cases and over 173,000 mortalities occurred in Africa (7). Almost all the regions across the world have recorded COVID-19 cases. However, some countries such as Turkmenistan have not yet registered any cases (8). The previous outbreaks of coronaviruses, including SARS and MERS, affected over 26 and 27 nations, (9) with morbidity of 8,000 and 2,519 for SARS and MERS, respectively (10). Compared to other pandemics, COVID-19 has caused higher morbidities and an overall lower case fatality ratio (CFR) as of 14^th^ July 2022 (11). Comorbidities such as diabetes, cancer, and hypertension have been linked to high mortalities due to COVID-19. On the other hand, Africa is known to be high in morbidities of infectious diseases such as HIV/AIDS and Malaria (12, 13, 14). Recent studies have been able to link such morbidities with COVID-19 co-infection and severe disease (15, 16).

SARS-CoV-2 has been conducted in countries such as South Korea, where in February 2020, nasopharyngeal (NP) and oropharyngeal (OP) samples were isolated from confirmed cases of COVID-19 patients where isolation and replication were confirmed through viral culture and gene sequencing (17). In June 2020, in the same country, serum, urine, and stool specimens were used to isolate the SARS-CoV-2, where the presence of the virus was evaluated using real-time RT-PCR (18). As of 15^th^ July 2022, the Kenya Ministry of Health had recorded a cumulative COVID-19 caseload of 336,445 and 5,668 related deaths (19). This paper describes the isolation, characterization, and viral dynamic of SARS-CoV2 Kenyan strain.

## Results

### Isolation of SARS-CoV-2 in culture

Three of the four clinical samples indicated cytopathic effect (CPE) in Vero CCL-81 cells (001, 003, and 004) three days post infection **(Figure 1 and 2)** following the initial infection and infectious culture fluids (ICFs) were frozen for further analysis. Sample 002 did not show CPE and ICF was kept for blind passaging. Sample 004 ICF was selected for sequencing.

**Fig. 1.**
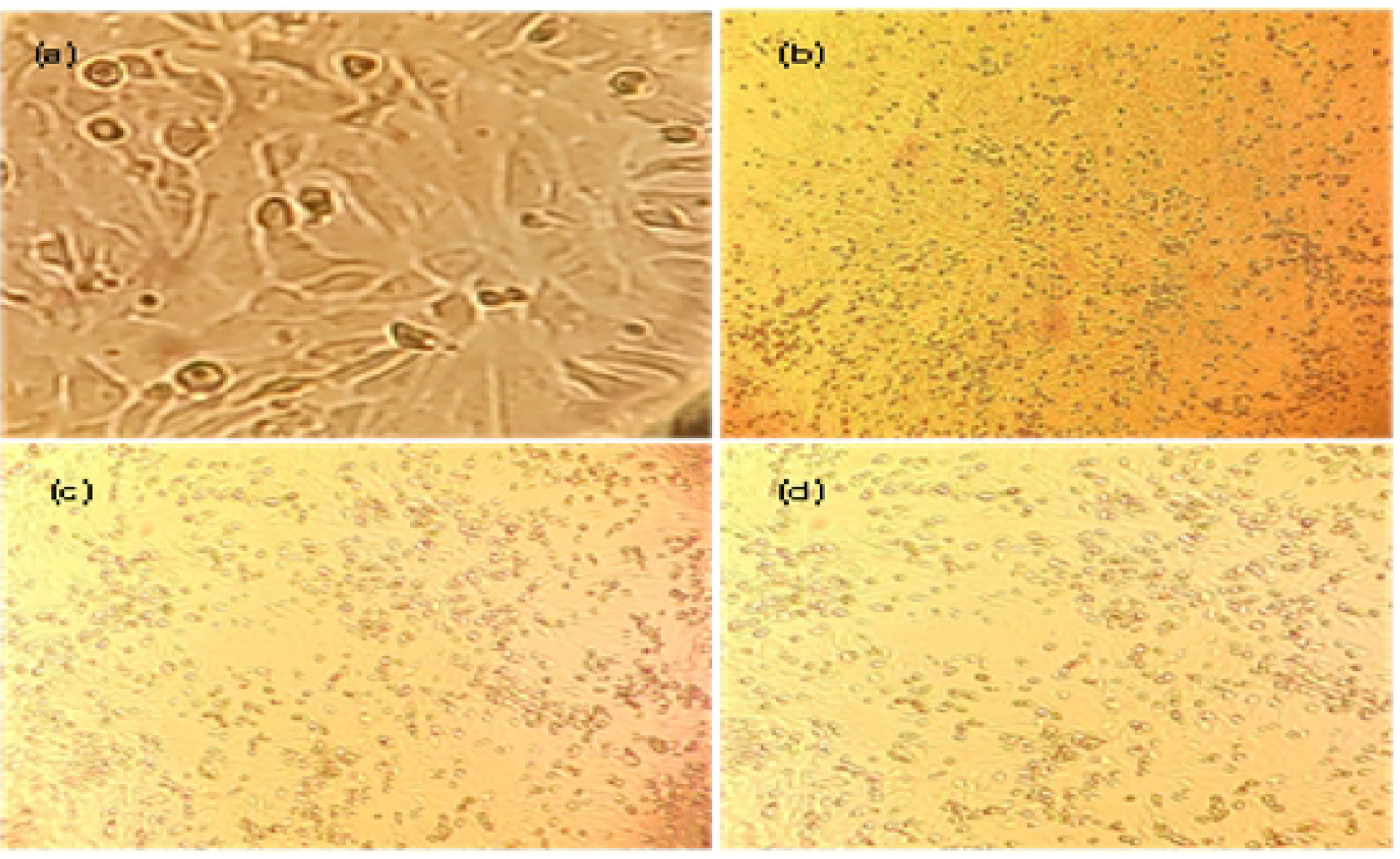
Phase contrast images of SARS-CoV-2 replication and cytopathic effects (CPE) in Vero CCL-81cell lines at 10X magnification. (a) Negative control cells inoculated with culture media, (b) Vero cells rounding up and floating in suspension 2 days post infection (c) Vero cells rounding up and floating in suspensions 3 days post infection(d) Cells rounding up 4 days post infection and CPE at 85%.

**Fig. 2.**
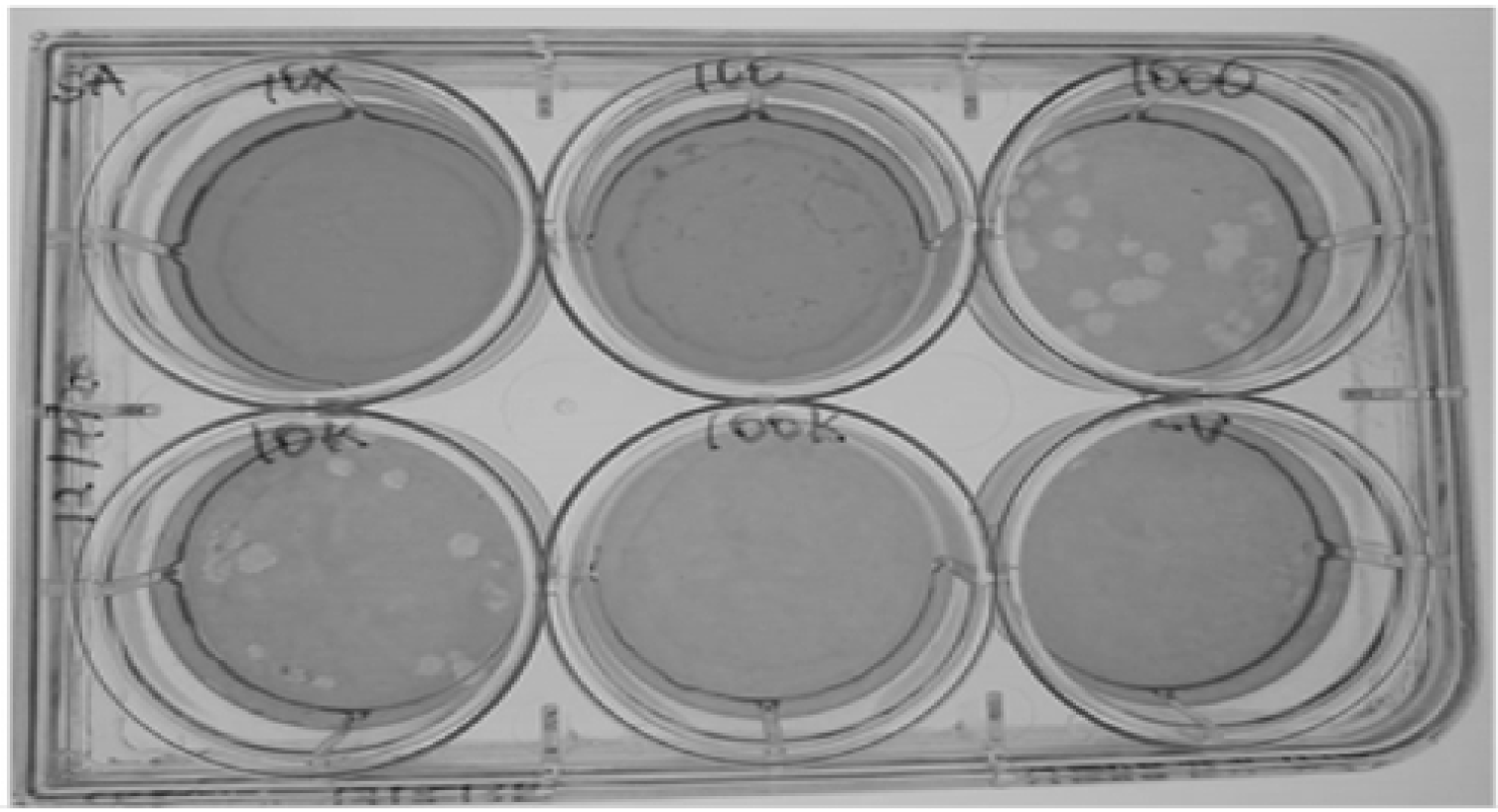
showing plaque formation of SARS-CoV-2 v passage 2 stocks at 3 days’ post infection on Vero cells CCL-81 cell lines.

### Quantification of SARS-CoV-2 by plaque assay

The isolated SARS-CoV-2 formed distinctly visible plaques, and the viral titer was determined as 1· 65 × 10^5^ pfu/ml.

### Conventional RT-PCR

A multiplex RT-PCR was conducted on 14 ICFs targeting the RdRp (344bp) and S (158bp) genes. Gel results were as shown below **(Figures 3 and 4)**.

**Fig. 3.**
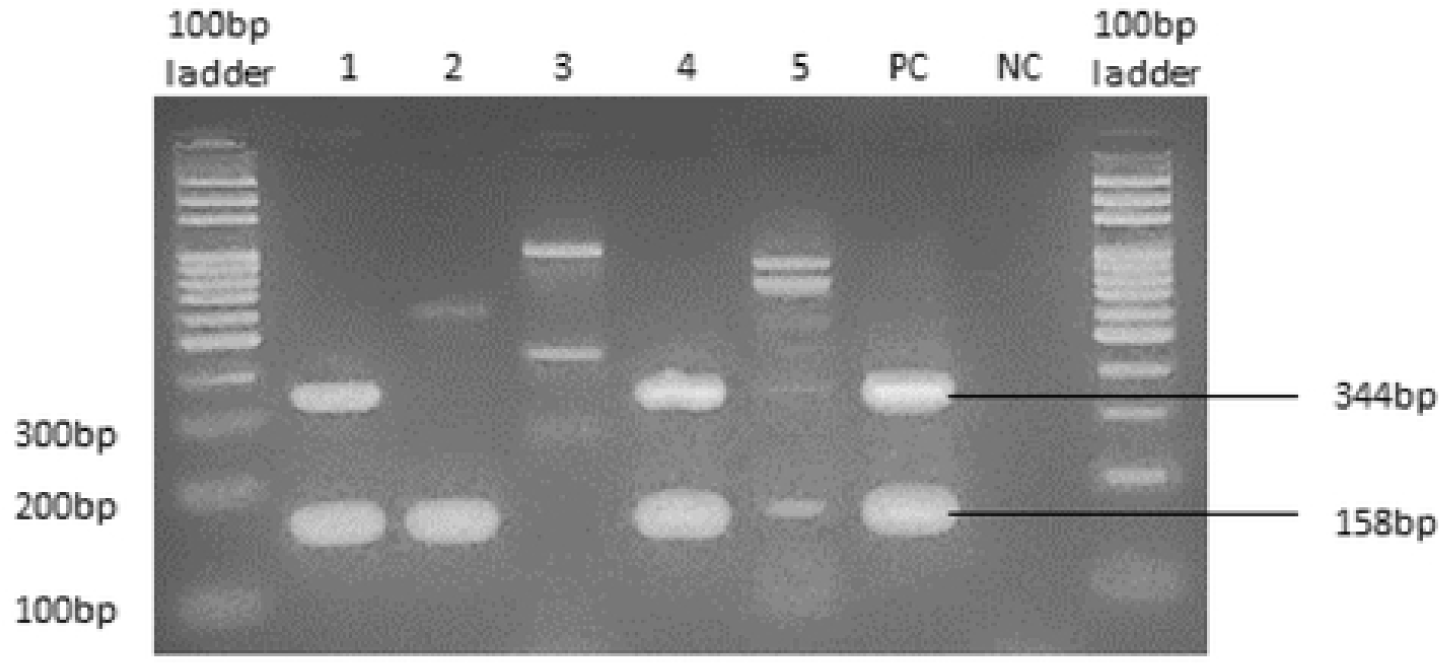
showing 5 PCR products run on 2% agarose gel. Clear bands targeting the RdRp and S genes can be seen for samples 1 and 4. Sample 2 was positive for the S gene PC jr.d NC are the positive and negative controls respectively.

**Fig. 4.**
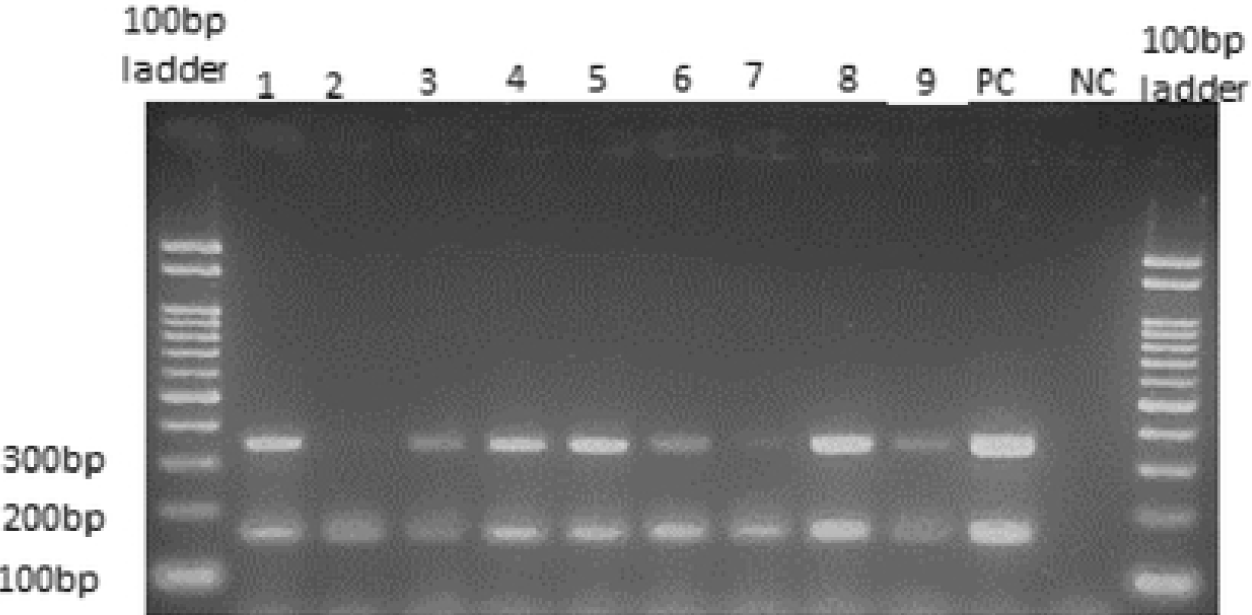
showing 9 PCR products run on 2% agarose gel. All saaples appeared positive far the S and RdRp genes.

Out of the 14 samples used in this study, 13 appeared positive for SARS-CoV-2 RdRp and S genes (92.9%).

### Relationship between the Local Isolates and the Global SARS-CoV-2

#### Whole-genome sequencing and phylogenetic analysis

We successfully assembled the whole genomes and confirmed the identity of SARS-CoV-2 virus isolates from our cultured isolates. The length of the assembled genomes from isolates 001 and 002 was 29,829 bp and 29,903bp, respectively. This represents 100% coverage of the reference genome. Comparisons of the sequences from this study to previous isolates from Kenya and the rest of the world in databases show apparent similarity, particularly to samples from Africa. We clustered the sequences based on ancestral similarity to confirm the clades where our isolates belong and further confirm the identity of our cultures. While our isolates clustered closely to other Kenya SARS-CoV-2 isolates, they did not fall entirely on the Kenyan clade but under a sub-clade comprising Benin isolates **(Figure 5)**. Further classification based on Next clade, analysis shows the strains to belong to the clade 20C, characterized by mutations [S:D80Y;N:S186Y;N:D377Y;ORF1a:T945I;ORF1a:T1567I;ORF1a:Q3346K;ORF1a:V3475F;O RF1a:M3862I;ORF1b:P255T;ORF7a:R80I] thought to have a Western USA origin **(Figure 6)**.

**Fig. 5.**
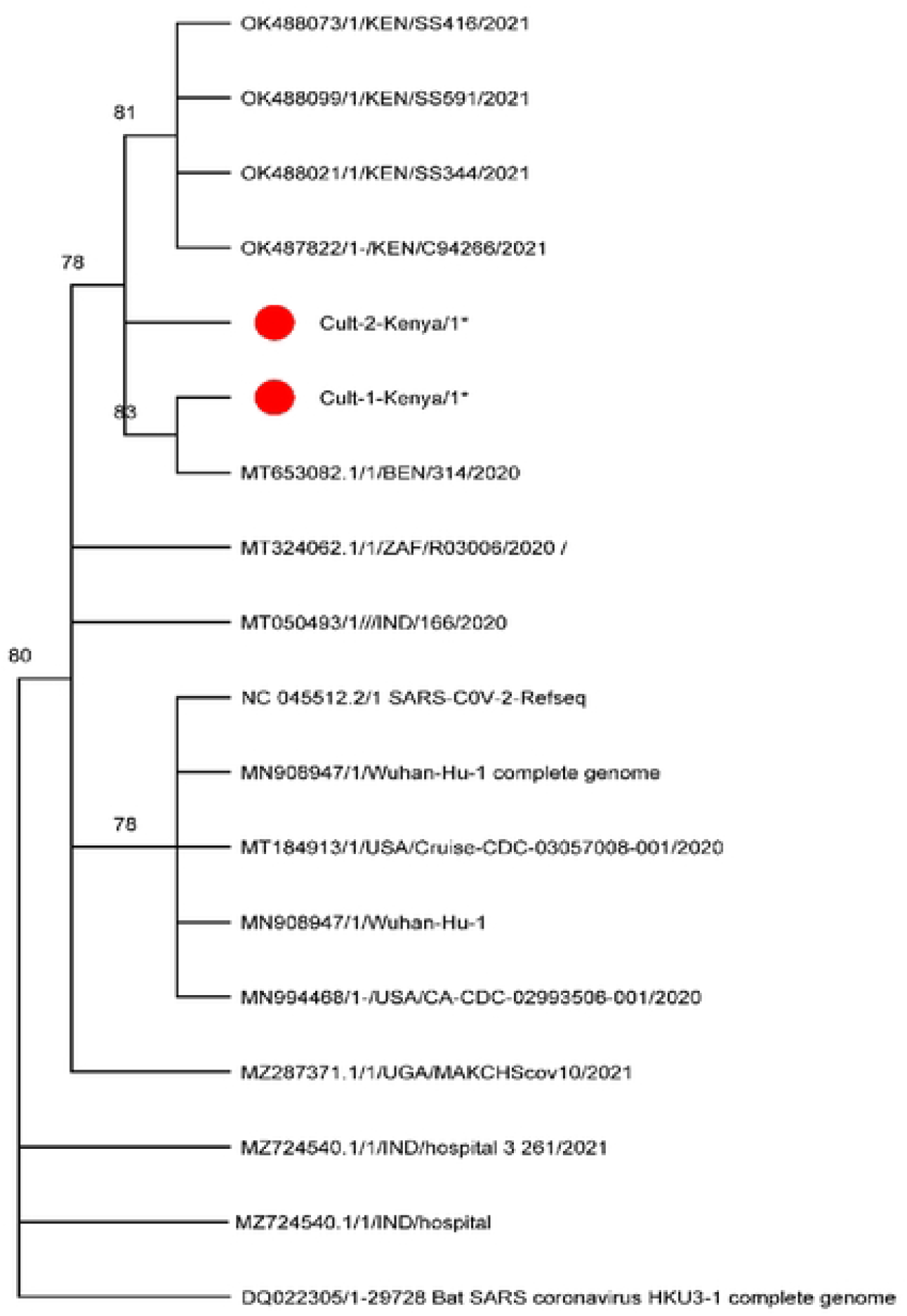
Phylogeny relationship between the two Kenya isolates and sequences from databases based on Neighbor-Joining parsimony. The percentage of replicate trees in which the associated taxa clustered together in the bootstrap test (1,000 replicates) are shown next to the branches. The MP tree was obtained using the Snbtree-Pruning-Regrafting (SPR) algorithm with search level 1, in which the initial trees were obtained by the random addition of sequences (10 replicates).

**Fig. 6.**
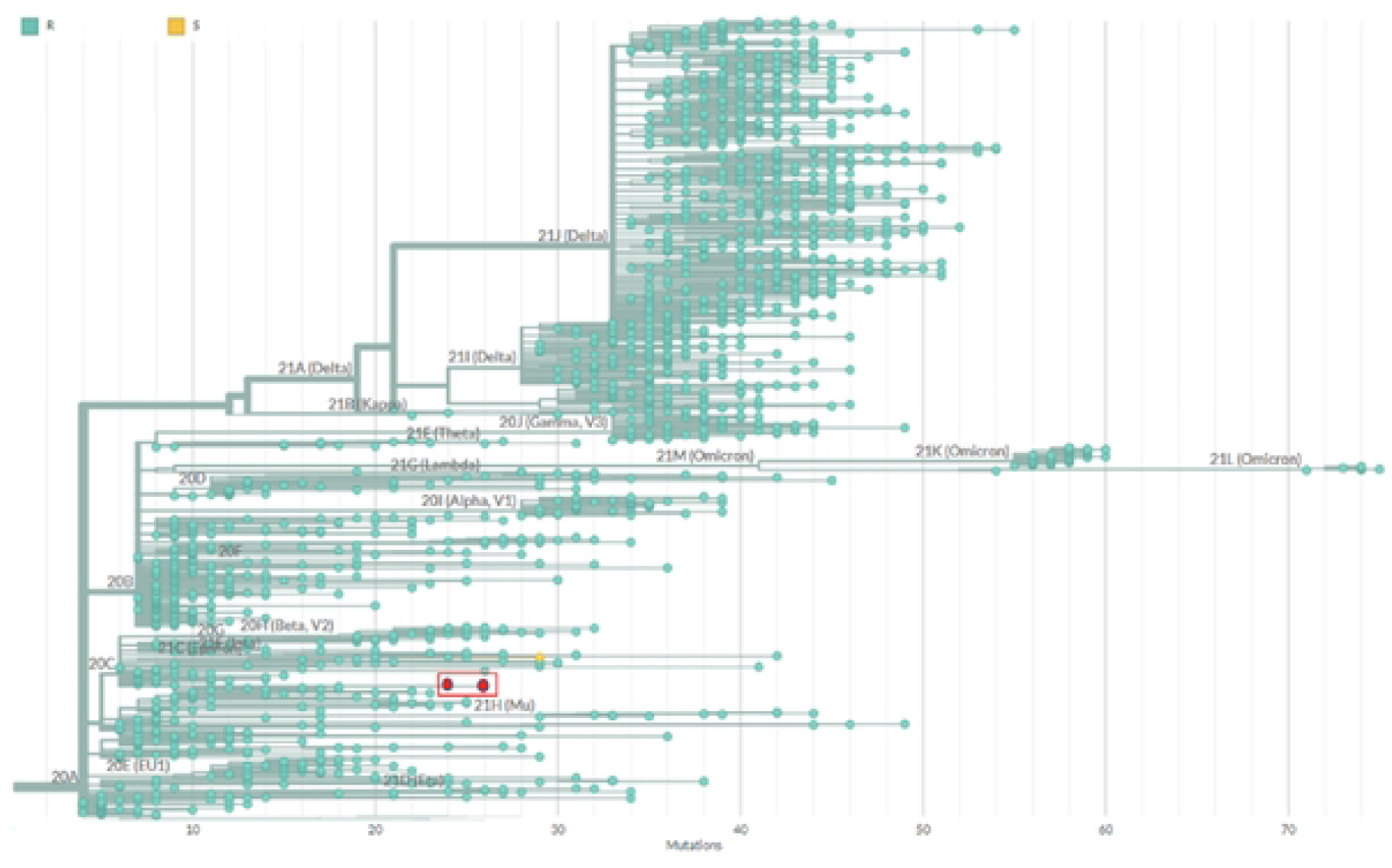
Phylogenetic tree built using Nextclade online software (https://clades.nextstrain.org/tree. (accessed on April 12^th^, 2022) and visualized using Auspice online tool (https://auspice.us(accessed on April 12^th^. 2022). Highlighted in red is the clade classification of sequences from this study.

#### Nucleotide substitutions and amino acid changes

To confirm substitutions at the protein level and follow up on the Nextclade classification of the isolates to the clade 20C, we aligned the sequences and profile mutations as shown in **Table 1**. While variations across the entire genome, pronounced differences in nucleotide sequences are evident in the S protein coding sequences (CDS) **(Supplementary Figure1)**. However, most of these are silent mutations, as we could identify only seven mutations in each sequence **(Table 1 and Supplementary figure 2)**.

**Table 1:**
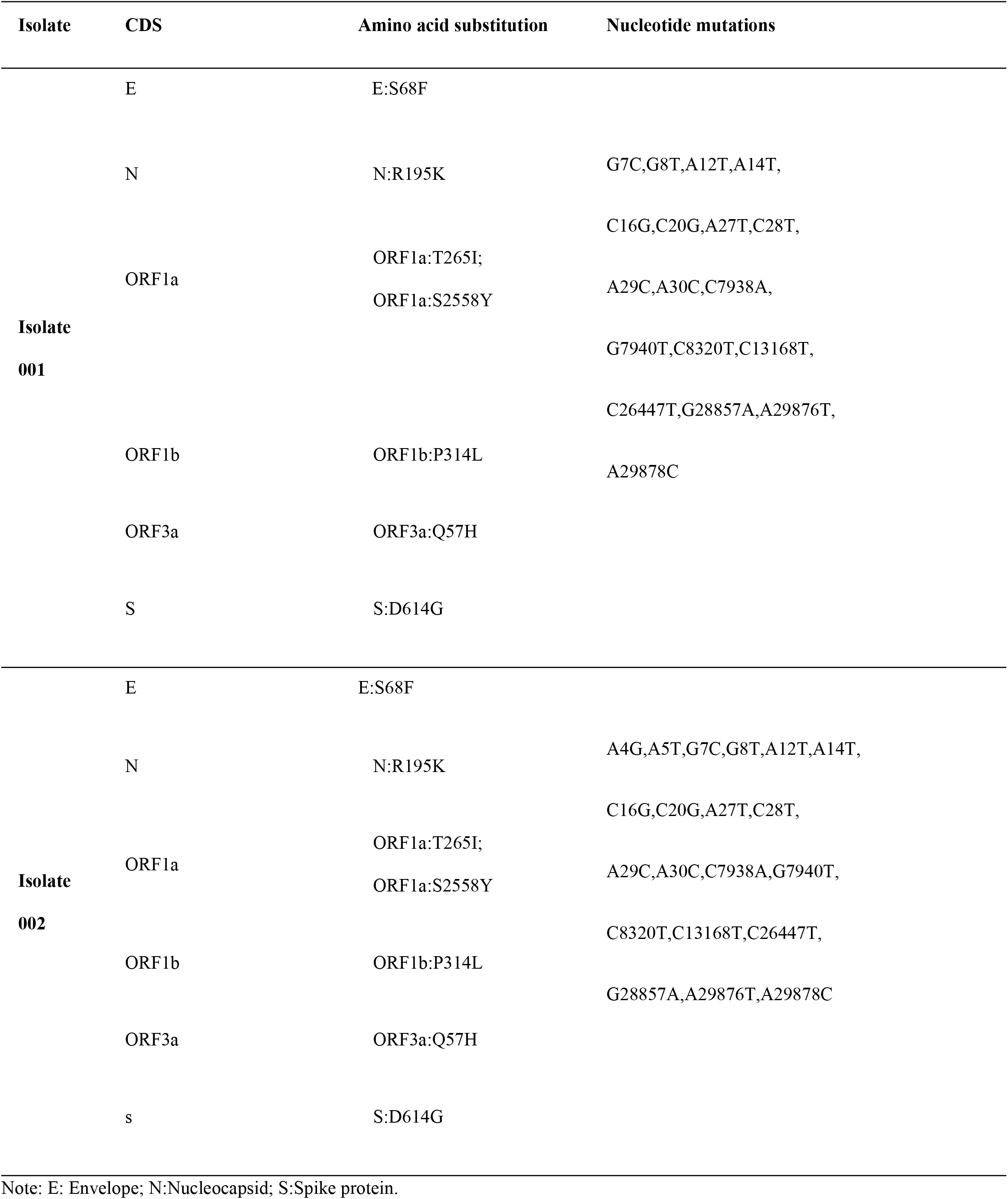
Mutations (substitutions) in amino acid sequences from the cultured isolates.

## Discussion

North America and Europe have shown higher COVID-19 CFR than Kenya and other African countries. This can be attributed to factors such as the high population of young people and the difference in climatic conditions (20). In our study, the SARS-CoV-2 virus was successfully isolated in Vero CCL-81 cells, reportedly successfully culturing the virus (21,22). NP and OP specimens of patients in Kenya were used for the virus isolation. The two different coronaviruses that occurred in Asia, SARS-CoV and MERS-CoV, were found to grow well on Vero CCL-81(23). In our analysis, SARS-CoV-2 replication was evident based on the observable CPE and subsequent plaque formation, implying the importance of the cells in-vitro study of the virus. A comparative study was done by Harcourt *et al*., (2020) (24), where they isolated and characterized SARS-CoV-2 from the first COVID-19 patient in the United States using both VeroE6 and CCL-81 cells. Both cells were observed to support the amplification and quantification of the virus. Our study is consistent with the Korean isolate, where the CPE occurred on Vero cell cultures after passaging on the third day (25). Isolation and characterization of SARS-CoV-2 will be helpful in studying immune response of COVID-19 patients to monitor ability to block viral infections. Furthermore, the local isolates can play a big role in antiviral research through locally available antiviral medications to treat or prevent COVID-19. Another area of much focus is in vaccine development as pharmaceutical companies and research institutions are in a race to develop robust vaccines that effectively end the pandemic. Therefore, as part of the global effort, research in the African continent such as Kenya to try to understand more about the virus pathogenesis is very essential for better future diagnostics and vaccine development.

After establishing an immense pattern of infection of SARS-CoV-2 on Vero CCL-81 cells, we extracted RNA from 14 ICFs and carried out conventional multiplex RT-PCR targeting the RdRp and S genes. RT-PCR has been considered to be the gold standard in SARS-CoV-2 identification; however, different testing kits give contrasting results based on specificity and sensitivity (26). The virus consists of the Spike (S), Membrane (M), Envelope (E), Nucleocapsid (N), and the RNA-dependent RNA polymerase (RdRp) proteins. Most of the RT-PCRs target the S gene due to its affinity and binding nature to the angiotensin converting enzyme 2 (ACE-2) receptors. Despite the positive outcome, previous studies have estimated the sensitivity of conventional RT-PCR to be between 70-98% and specificity approximated at 95% (27, 28). It was also shown that the diversity of SARS-CoV-2 can affect the outcome of an RT-PCR analysis (29). Our study was successful in targeting both genes showing they can be effective in detecting the virus from clinical samples. SARS-CoV-2 continues to affect countries of the world. Studying in detail the virus genome will open up more features in tracing the virus source path and source of infection (30). In this study, several genomes were aligned together with our two isolates for characterization based on phylogenetic analysis. The results show that our two isolates share a close relationship with Benin isolate while the Nextclade analysis showed direct linkage to the Western U.S.A isolate. The nucleotide sequences and their resultant amino acid (AA) composition were further investigated for any possible mutations. Most of the SARS-CoV-2 mutation studies have been done along with the S and N proteins. D614G mutation along the S protein has been reported from previous analysis, seeming to be a predominant one (31,32), enhancing the binding affinity of SARS-CoV-2 to the ACE2 receptors (33). Both isolates had this mutation at the S protein, with recent studies showing the AA change to aid in faster viral transmission (34). All nucleotide substitutions were common in the isolates across the different genes despite most of them giving silent mutations. An R195K AA change in the N protein was reported in this study though from a different position from the one found in this study (R203K) (35), which may be significant in blocking an immune response through antibody production among infected patients. C-T single nucleotide polymorphism (SNP) was seen to occur along the open reading frame-1a (ORF1a). Previous studies have shown that the C-T SNP at position 28144 led to a Serine-Leucine amino acid change, thereby elucidating a higher transmission rate (36). In our study, a T265I and S2558Y AA substitutions were found to occur along the ORF1a.

So far, a number of variants with multiple mutations have been discovered since the emergence of the pandemic. This shows that the SARS-CoV-2 virus continues to evolve so as to evade the host defense system despite efforts to eliminate it through the use of currently developed vaccines and antivirals. Our study was successful in deciphering the evolution of the SARS-CoV-2 within the Kenyan population through phylogeny and mutational analysis.

## Conclusion

This study showed that Vero CCL-81 cells could support SARS-CoV-2 virus isolation and characterization through the formation of viral plaques. Detection of SARS-CoV-2 targeting the S and RdRp genes was effective. However, a limitation of this study was that the isolates cultured were from samples collected in July, 2020, when the pandemic emerged, with very few strains reported. As we are aware, the virus has dynamically evolved; hence future studies should focus on recent samples to sequence viral genomes from different isolates collected in different regions to increase the probability of identifying multiple strains circulating in Kenya and even possible novel mutations, which will also offer a road map in the development of diagnostics and vaccines to control the spread of the disease. Another significant area of study could be on co-infections of SARS-CoV-2 with other endemic diseases such as HIV/AIDS and malaria that have continued to affect the Kenyan population for a long time so as to depict disease severity due to co-infection within a human host.

## Materials and Methods

### Ethical Statement

The Scientific and Ethics Review Unit (SERU-KEMRI) approved this study under protocol number KEMRI/SERU/CBRD/210/4027.

### Specimen collection

NP and OP swabs were collected as described (37) from symptomatic patients reported to health facilities in Kenya in July 2020, and placed in a virus transport medium (VTM). To maintain confidentiality, the samples were assigned unique identities and delinked from the patient information. The samples were transported to KEMRI in cold packs and stored at the Sample Management and Receiving Facility (SMRF) at -80°C pending processing. The handling and processing of the samples were done at the KEMRI biosafety level 3 laboratory, where analysis was performed as described in the US Centers for Disease Control (CDC) guidelines (38).

### Isolation of SARS-CoV-2 from clinical samples

Isolation and propagation of high titers of infectious SARS-CoV-2 were done in a biosafety level 3 laboratory by qualified staff using a modified method by Tastan and colleagues (39). Following confirmation of the virus by conventional RT-PCR, low passage Vero CCL-81 cells were cultured in from the cell bank were thawed into Minimum Essential Medium (MEM, Life Technologies Carlsbad, CA) supplemented with 10% (v/v) fetal bovine serum (FBS) (Gibco) and 2% Penicillin-Streptomycin-Amphotericin solution (Gibco). The frozen samples were thawed on ice, and a 10x dilution was prepared in single-strength MEM. A 100 μl of the diluted virus was inoculated into wells of a 96-well culture plate of Vero CCL-81 cells. Upon confirmation of CPE under an inverted microscope, the contents of the wells were sequentially propagated in culture plates and flasks (NUNC, Roskilde, Denmark) containing 90% confluent monolayers of Vero cells. In all cases, plates and flasks were maintained in a humidified 5% CO_2_ atmosphere in an incubator at 37° C. Cells were monitored daily for CPE and at 85% of CPE; the ICF from the 75cm^2^ flask was harvested by centrifuging for 10 minutes at 4° C. The supernatant was aliquoted and kept at -80° C as seed virus stock.

### SARS-CoV-2 viral titer determination

Plaque assays were done based on SARS-CoV-2 and MERS-CoV protocols (40) with a few modifications. Vero CCL-81 cells were seeded in 6-well culture plates to attain a 90% cell confluence before inoculation with 200μl of seed virus stock. After overlaying the infected cells with agarose gel, the plates were incubated at 37°C in a 5% carbon dioxide gas atmosphere for 3-4 days until plaques were evident. Staining was done with neutral red dye to visualize the plaques, and after overnight incubation, the plaques were counted and determined using the formula, Titer (pfu/ml) = number of plaques X dilution factor X 1/volume of virus added to cells in a well (ml).

### Viral RNA Extraction

Viral RNA was extracted from the supernatants using the viral RNA mini kit (QIAGEN, Hilden, Germany) according to the manufacturer’s instructions and temporarily kept at -20° C awaiting reverse transcription.

### Reverse-transcription Polymerase Chain Reaction (RT-PCR)

Reverse transcription polymerase chain reaction was done as described (19). Primers targeting the RdRp gene forward (5’-CAAGTGGGGTAAGGCTAGACTTT-3’) and reverse (5’– ACTTAGGATAATCCCAACCCAT-3’) with an amplicon size of 344bp and S gene forward (5’-CCTACTAAATTAAATGATCTCTGCTTTACT-3’) and reverse (5’-CAAGCTATAACGCAGCCTGTA-3’) with an amplicon size of 158bp were used in this assay (41). Reverse-transcription was carried out at 50 °C for 30 minutes followed by 95 °C for 15 minutes, 50 cycles of 95 °C for 60s, 55 °C for 60s and 1 step at 72 °C for 10 minutes. Positive and negative controls were included in all assays. For the no template control, 5 μl of nuclease-free water was added in place of the RNA template. Likewise, 5 μl of RNA with a known Ct value was added in place of template RNA to serve as the positive control. Cycles were run on Applied Biosystems ABI 7500 Fast instrument. Finally, gel electrophoresis was conducted using 2% agarose gel at 100V for 45 minutes, stained for one hour and visualized under UV-transilluminator (Maestrogen Inc, Taiwan).

### Whole genome sequencing of the RT-PCR Product

A total of 50ng cleaned-up PCR product was repaired using NEBNext Ultra II End repair/dA-tailing reagent and barcoded using NEBNext Ultra II Ligation Master Mix (New England Biolabs Inc. Massachusets, USA), following manufacturer’s guidelines. The barcoded DNA was pooled and purified using AMPure XP beads (Beckman Coulter Life Sciences, Indianapolis, USA) and quantified using fluorometer (DeNovix Inc. Wilmington, USA). At least 2ng/ul of the DNA libraries were loaded on a flow cell and sequenced on a MinION (Oxford Nanopore Technologies, Oxford, United Kingdom), which sequences long reads of the ssDNA molecule with cartridges containing 2048 nanopores arranged in 512 channels. Leader and hairpin adapters were used to prepare dsDNA that was recognized by specific signal generated by the apurinic and /or apyrimidinic sites in the hairpin. Sequence consensus was built following a successful reading of the DNA strands.

Raw read sequences from the Oxford Nanopore MinION sequencing platform were evaluated for quality based on FastQc reports and deposited in the NCBI SRA database with accession PRJNA825709. The quality threshold was set at 20, and a minimum length of 150 was allowed. Trimmomatic tool (42) was used to remove adapter sequences, followed by a post-trimming quality assessment based on the FastQC report. We used SPADES (43) to assemble raw sequences with reference genome retrieved from the SARS-CoV-2 RefSeq database: https://ftp.ncbi.nlm.nih.gov/refseq/release/viral/; March 10^th^ version. Raw sequences were deposited in the NCBI’s Sequence Retrieval Archive (SRA) under accessions SRS12585595 and SRS12585594. The sequences were identified using nucleotide Basic Local Alignment Search Tool_BLAST (44). Upon confirming the sequence, we performed cluster analysis of the isolates using Nextstrain/Nextclade v1.14.0 (45) server. Equally, mutations in amino acid residues were conducted using Nextclade. Sequence alignment and phylogenetic comparisons were achieved using MUSCLE (46) and MEGAX (47).

## Acknowledgments

We sincerely thank KEMRI for their immense support throughout the research process and the Internal Research Grants of KEMRI which made the work successful. This manuscript has been written with the permission of the Director General and Chief Executive Officer, KEMRI.

